# Rice LEAFY COTYLEDON1 hinders photosynthesis in the embryo development to promote seed dormancy

**DOI:** 10.1101/2021.08.18.456739

**Authors:** Fu Guo, Peijing Zhang, Yan Wu, Guiwei Lian, Wu Liu, B Buerte, Chun Zhou, Ning Han, Muyuan Zhu, Lin Xu, Ming Chen, Hongwu Bian

**Author notes:** Corresponding author: Hongwu Bian, Ming Chen, Lin Xu. These authors contributed equally to this work (Fu Guo, Peijing Zhang, Yan Wu). **Author contributions:** FG, HB, MC, and LX designed the research; PZ and YW performed the data analyses; GL, WL, B, and CZ performed the experiments; FG and HB wrote the manuscript, NH, MZ, MC, and LX revised the manuscript. FG, PZ, and YW contributed equally to this work. **Competing interests:** The authors declare no competing interests.

## Abstract

LEAFY COTYLEDON1 (LEC1) is the central regulator of seed development. During seed development, rice embryo photosynthesis is completely blocked, which is different from Arabidopsis green embryo. However, effects of LEC1 on photosynthesis in developing seeds is largely elusive. We generated OsLEC1 mutants using the CRISPR/Cas9 technique. *Oslec1* mutant seeds lost the ability of dormancy and triggered photosynthesis in embryos at the early developing stage. Transcriptome analysis demonstrated that *Oslec1* mutation promoted photosynthesis and altered diverse hormonal pathways and stress response contributing to seed dormancy. Further, genome-wide identification of OsLEC1 binding sites demonstrated that OsLEC1 directly bound to genes involved in photosynthesis, photomorphogenesis, as well as abscisic acid (ABA) and gibberellin (GA) pathways, in seed maturation. We illustrated an OsLEC1-controlling gene network during seed development, including the interconnection between photosynthesis and ABA/GA biosynthesis/signalling. Our findings suggested that OsLEC1 is an inhibitor of photosynthesis during embryo development to promote rice seed maturation. This study would provide new understanding for the OsLEC1 regulatory mechanisms on photosynthesis in the monocot seed development.

## Introduction

Seed dormancy plays an important role in agriculture. It is a characteristic of the seed that determines the conditions required for germination (1, 2). Extremely strong dormancy leads to a low germination rate and irregular emergence of seedlings, has an impact on sowing time, and can even affect the final yield. On the contrary, inappropriate loss or release of seed dormancy results in the rapid germination of freshly matured seeds or even pre-harvest sprouting (PHS), which has become an important factor restricting the yield of cereal crops (3). Seed dormancy and germination has been studied extensively in the past; however, the co-regulation of transcription factors and the regulatory network involved in seed dormancy remains largely unknown in monocot crops. Seed dormancy has been called “one of the most mysterious phenomena in seed biology” (4).

Plant seed development is divided into two phases: morphogenesis phase and maturation phase. During the morphogenesis phase, the basic body plan of the embryo and endosperm are established (5, 6); chloroplast biogenesis and photosynthesis are also initiated during this period in many angiosperm taxa (7). During the maturation phase, the accumulation of storage compounds, and the induction of desiccation tolerance and preparation for seed dormancy are the main processes (8).

Seed development is genetically controlled by at least four genes, leafy cotyledon 1 (*LEC1*), *LEC2, FUSCA 3* (*FUS3*), and abscisic acid insensitive 3 (*ABI3*) (9-12), and *lec1, lec2, fus3*, and *abi3* mutants represent the reduction of desiccation tolerance and seed dormancy (11-19). Such regulators interact in a network to induce seed dormancy (20). In addition, it is widely recognised that phytohormone abscisic acid (ABA) and gibberellin (GA) are the primary hormones regulating seed dormancy and germination (21-24). LEC1 acts as a central regulator of seed development through combinatorial binding with aba-responsive element-binding protein 3 (AREB3), basic leucine zipper 67 (bZIP67), and ABI3 (8).

In Arabidopsis, loss-of-function *lec1* mutations cause defects in storage proteins and lipid accumulation, acquisition of desiccation tolerance, and suppression of germination and leaf primordia initiation (12, 15, 19, 25). Moreover, ectopic expression of *LEC1* induces embryonic development and the activation of genes involved in maturation and storage, as well as lipid accumulation in vegetative organs (13, 26, 27). Few studies have explored the effects of the overexpression of *LEC1* in carrot, maize, and rice. Expressing carrot *C-LEC1* driven by the Arabidopsis *LEC1* promoter could complement the viviparous and desiccation intolerant defects of Arabidopsis *lec1-1* mutant (28). Overexpression of maize (*Zea mays*) *leafy cotyledon 1* (*ZmLEC1)* increases seed oil production by up to 48% but reduces seed germination and leaf growth (29). In rice, *OsLEC1* overexpression results in abnormalities in the development of leaves, panicles, and spikelets (30). OsNF-YB9 and OsNF-YB7 (OsLEC1) are homologous to Arabidopsis LEC1. Heterologous expression of either OsNF-YB9 or OsNF-YB7 in Arabidopsis *lec1-1* complements the *lec1-1* defects. OsNF-YB7 defect causes lethality, and loss of OsNF-YB9 function causes abnormal seed development (31). Recent transcriptional studies showed that LEC1 could promote but is not absolutely required for photosynthesis and chloroplast biogenesis in Arabidopsis and soybean seed development (18, 32). However, how LEC1 regulates chloroplast biogenesis and photosynthesis during seed development is not clear.

In the present study, we generated *OsLEC1* mutants using a gene-editing technique, which is an orthologue of Arabidopsis *LEC1*, acting as the central regulator of seed development. Phenotype analysis revealed that dry seeds of *Oslec1* mutants could not germinate but fresh seeds of the mutants germinated normally, suggesting the absence of dormancy. Notably, embryos of mutants turned green during early seed development. Subsequently, RNA-seq transcriptional profiling and ChIP-seq were used to identify the underlying mechanisms via which *OsLEC1* directly regulates photosynthesis and seed dormancy during rice seed maturation. Our studies provide new evidence that OsLEC1 is an inhibitor of photosynthesis during rice embryo development, contributing to seed dormancy.

## Results

### OsLEC1 deficiency disrupts seed dormancy and germination

We first established that LEC1 was largely conserved while typically different in a few sites between monocots and dicots by performing multiple alignments (Supplemental Figure 1A, B). To determine the physiological functions of OsLEC1 in rice, we used the CRISPR/Cas9 technology to knock out *OsLEC1*. We generated more than 30 transformant plants for the T0 generation, in which seven homozygous deletion mutants were screened using PCR and DNA sequencing (data not shown). DNA sequencing analysis showed that *Oslec1-1* and *Oslec1-2* contained a 1 bp deletion (G) at the gRNA1 site, and a 1 bp insertion (A or T) at the gRNA2 site (Figure 1A), leading to frameshift mutations and generating premature stop codons at aa 157 (Supplemental Figure. 1C). Thus, *Oslec1-1* and *Oslec1-2* mutations disrupted the original protein structure from aa 254 to 157. Except for the shorter height of the mutants than the wild type plant, we did not observe obvious abnormal morphological phenotypes including the tillering or grain number of the mutants (Supplemental Figure 2). When >90% of the wild type seeds germinated after drying at 37°C, none of the mutant seeds germinated (Figure 1B, 1D). Surprisingly, freshly harvested mutant seeds germinated and even rooted 24 h after imbibition, when the wild type plant had not germinated yet (Figure 1E, Supplemental Figure. 3). Finally, >90% of *Oslec1-1* and *Oslec1-2* fresh seeds germinated, similar to the wild type plant (Figure 1C). Further, 2,3,5-Triphenyltetrazolium chloride (TTC) staining results, showed that the dry seeds of *Oslec1* mutants displayed green primordium without any red tissues, indicating completely lethal embryos in the dry mutant seeds (Figure 1D). Interestingly, green leaf primordia were observed in *Oslec1-1* and *Oslec1-2* embryos but not in wild type ones, suggesting chlorophyll accumulation in the *Oslec1* embryo. Meanwhile, we tested the embryo vitality of freshly harvested seeds (about 25 days after pollination [DAP]) using TTC staining. The embryo vitality of *Oslec1* fresh seeds was similar to that of the wild type ones (Figure 1E).

**Figure 1.**
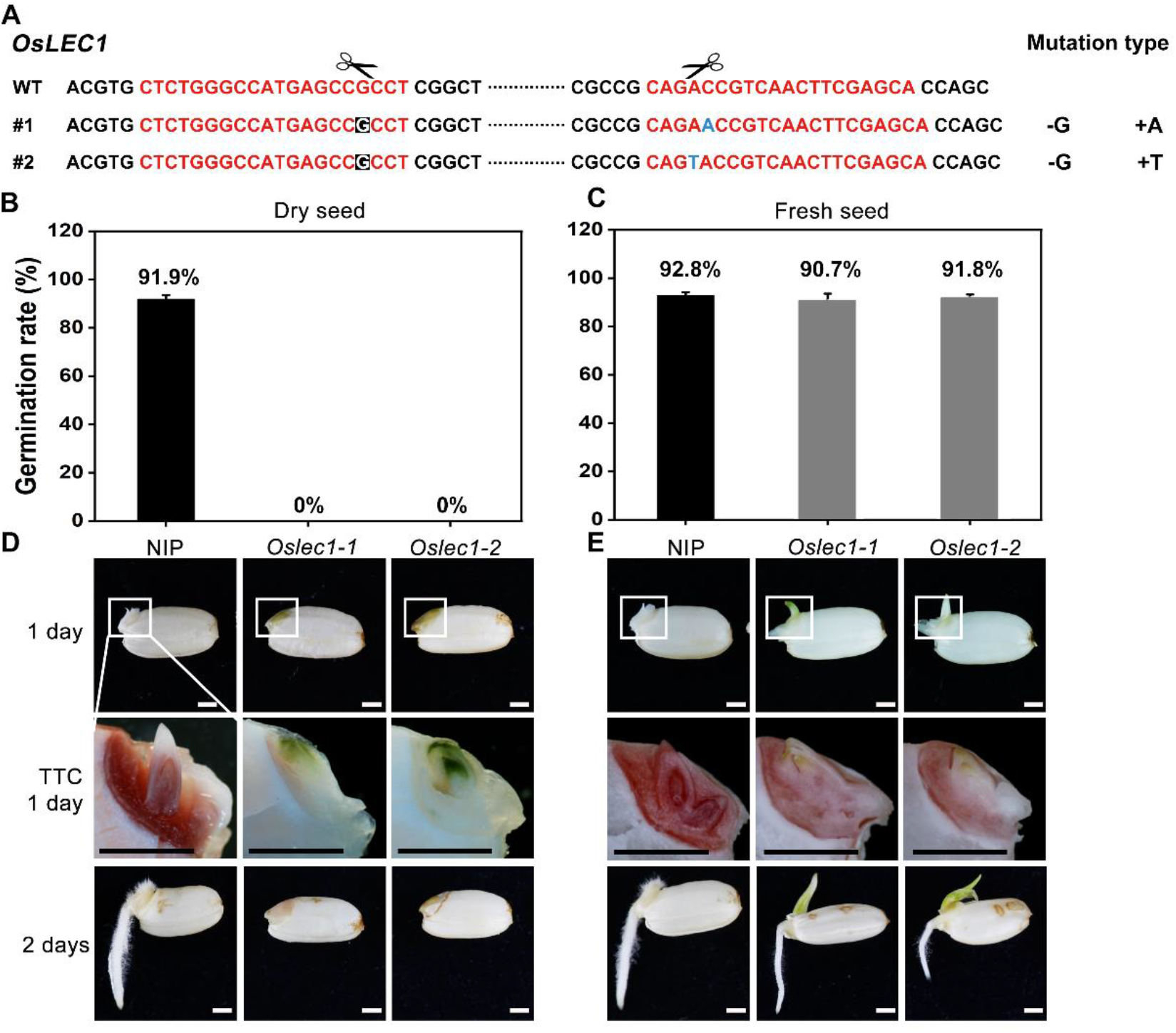
Germination phenotypes of *Oslec1* mutants. (**A)** Mutation sites in the *Oslec1 mutant* DNA. The red sequence represents the designed guide RNA (gRNA) for the wild type rice and the scissors represent the predicted knockout sites. The letters with a black background represent the deleted bases, and the blue letter represents the inserted base. (**B-C)** Germination rates of dry seeds and the fresh seed of the wild type and *Oslec1* mutants after 1 day of imbibition at 37°C. **(D-E)** Photos of the wild type and *Oslec1* mutant seeds after 1 day of imbibition (1 day) and 1 day on wet filter paper (2 days). Photos of group ‘TTC 1 day’ were the amplification of the regions in white boxes in group ‘1 day’, indicating the embryos stained by TTC for 3 h at 37°C. Scale bars=1 mm.

Our results revealed that the *Oslec1* mutants disrupted desiccation tolerance but initiated chlorophyll accumulation in seed development, suggesting that *OsLEC1* acts as a key regulator of seed dormancy in rice.

### OsLEC1 deficiency leads to a green embryo during seed development

Further, we examined the embryo phenotypes of the mutants and wild type plants at different stages of rice seed development. In contrast to the wild type ones, *Oslec1* mutant embryos displayed green apical shoots 7 DAP (Figure 2A). Subsequently, the apical shoot was green in *Oslec1* mutant embryos 25 DAP (Figure 2B). Surprisingly, after being imbibed in water for only 3 h at 37°C, the epiblasts covering the embryos were split by the sprouting shoots in the *Oslec1* mutants, but not in the wild type plants (Figure 2C). *OsLEC1* deficiency led to a green embryo during seed development, indicating the mutants completely lost dormancy.

**Figure 2.**
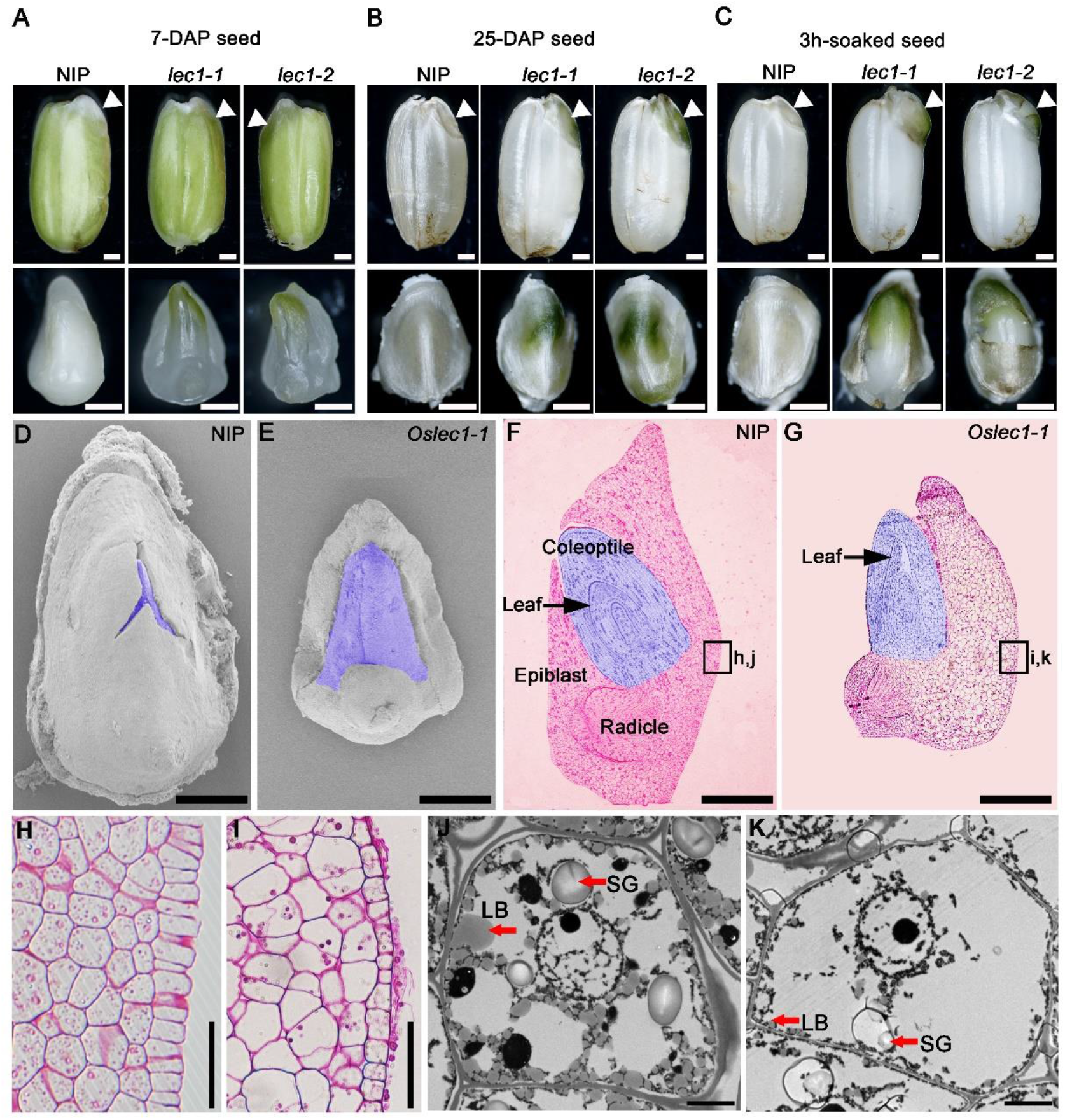
Phenotypes of the developing seeds of *Oslec1* mutants. **(A)** Seeds and embryos (7-DAG) of wild type plants and *Oslec1* mutants. **(B)** Seeds and embryos (25-DAP) of wild type plants and *Oslec1* mutants. **(C)** Seeds and embryos of wild type plants and *Oslec1* mutants were soaked in water for 3 h at 37°C. Scale bars=500 µm. White triangles indicate the embryonic regions in complete seeds. **(D-E)** Scanning electron micrographs of embryos (7-DAP) of wild type plants and *Oslec1* mutants. The purple-coloured regions indicate the germs of wild type plants and *Oslec1* mutants. Bars=500 µm. **(F-G)** Longitudinal resin sections of embryos (7-DAP) of wild type plants and *Oslec1* mutants. The purple-coloured regions indicate the germs of wild type plants and *Oslec1* mutants. Black arrows indicate the leaves in the coleoptiles. The leaves of *Oslec1* mutants are stretched out, and the epiblast wrapping the coleoptile is missing. Scale bars=500 µm. **(H-I)** Amplification of the regions in black boxes in f and g, showing the difference in scutella parenchyma between the wild type plants and *Oslec1-1* mutants. Scale bars=50 µm. **(J-K)** Electron micrographs of wild type and *Oslec1-1* embryos. Wild type (J) and *Oslec1-1* (K) scutellum sections from 7-DAP embryos. Scale bars=2 µm.

Scanning electron microscopy (SEM) showed that 7 DAP, the shoot apical meristems in wild type embryos were embedded in the covering epiblast (Figure 2D), while those in the mutants were exposed (Figure 2E). Semi-thin longitudinal sections of embryos showed an ordered three-layer leaf primordium in the intact coleoptile of wild type embryos (Figure 2F), while in *Oslec1* mutant embryos 7 DAP, the primary leaves were spread out (Figure 2G) and the scutella parenchyma was abnormal compared with those of the wild type plants (Figure 2J, 2K). Transmission electron microscopy (TEM) showed many lipid bodies and large starch grains in wild type embryos 7 DAP, while major storage products were almost absent in *Oslec1* mutant embryos (Figure 2H, 2I). Thus, the green embryos of *Oslec1* mutants showed that photosynthesis, or light response, and leaf primordia initiation were triggered in the embryo. The above data suggested that *OsLEC1* might play a crucial role in the negative regulation of photosynthesis, early germination, and accumulation of storage compounds in rice seed development.

### *OsLEC1 is* expressed predominantly in immature embryos during early seed development

We analysed the expression pattern of *OsLEC1* based on an expression database (http://expression.ic4r.org/) (33) (Supplemental Figure 4A). The results showed that *OsLEC1* was highly expressed in the callus and immature embryos. Subcellular localisation showed that OsLEC1-GFP was enriched in the nucleus, which was indicated by OsIAA1-mCherry in rice protoplasts (Supplemental Figure 4B).

*pOsLEC1:GUS* was expressed strongly in the dorsal section of immature embryos within 4-7 DAP (Figure 3A, 3B) and then declined to a relatively low level at 25 DAP (Figure 3C-E). Further, semi-thin sections of resin-embedded embryos showed that *OsLEC1* was predominantly expressed in the cells of scutella parenchyma, especially in the apical and basal part of the embryo 8 DAP (Figure 3G-I). Expression patterns of *OsLEC1* suggested its function in the early stage of rice embryo development.

**Figure 3.**
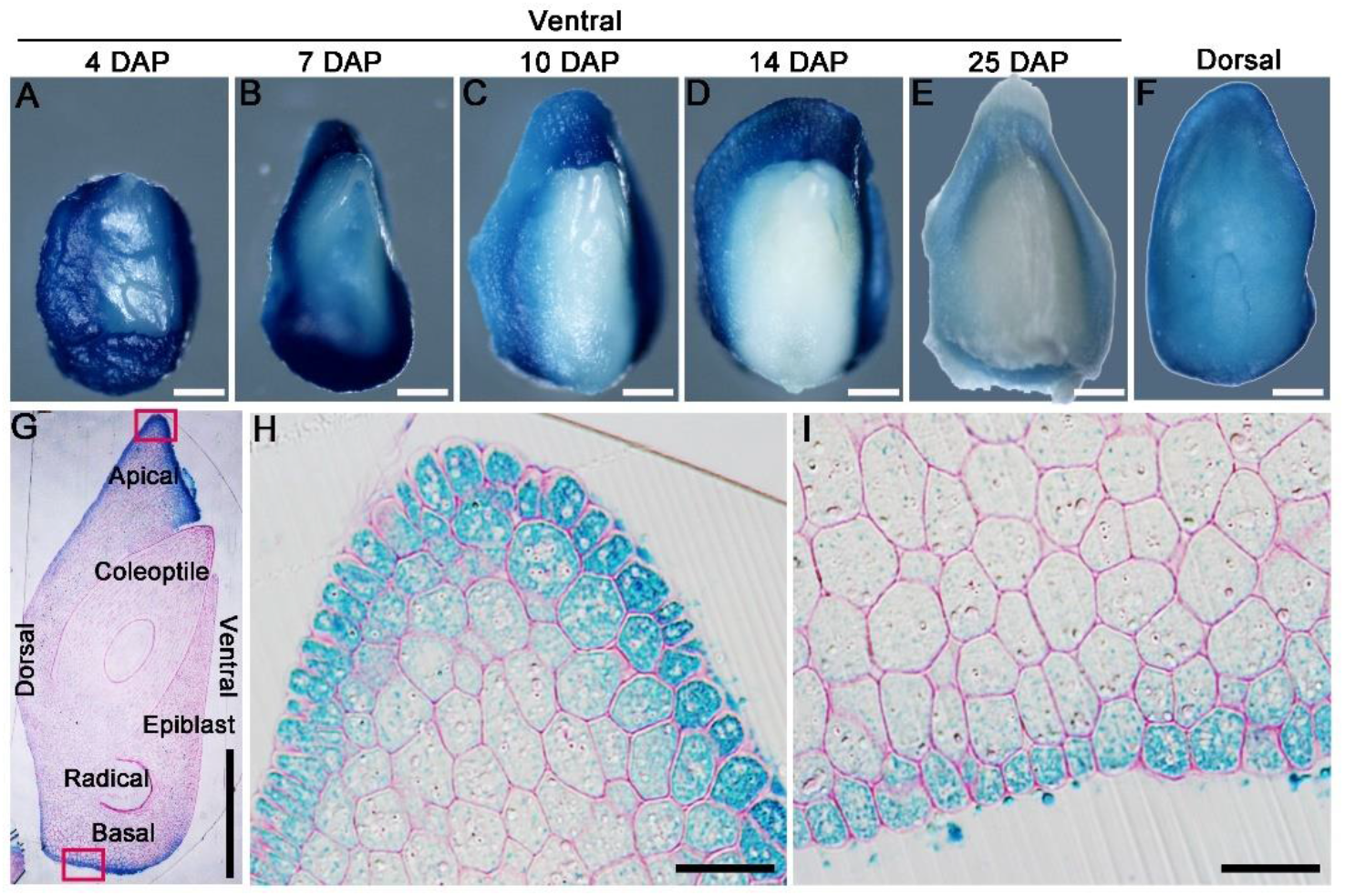
Expression pattern of *OsLEC1*. **(A-F)** Expression of *OsLEC1* in developing embryos. **(A-E)** Ventral side of *pOsLEC1:GUS* embryos (4–25 DAP) stained in X-Gluc solution for 3 h at 37°C. Scale bars=200 µm. **(F)** Dorsal side of *pOsLEC1:GUS* embryos (14 DAP) stained in X-Gluc solution for 3 h at 37°C. Scale bars=200 µm. **(G)** Longitudinal resin section of embryos (7 DAP) of *pOsLEC1:GUS* embryo stained in X-Gluc solution overnight at 37°C. Scale bars=200 µm. **(H-I)** Amplification of the regions in black boxes in g, showing the apical (**H**) and basal (**I**) parts of the scutella parenchyma, respectively. Scale bars=50 µm.

### OsLEC1 mutation influences multiple developmental processes

To identify the genes that were controlled or regulated by *OsLEC1*, we performed RNA-seq in *Oslec1-1* mutant and wild type embryos at two different stages of seed development: the early-stage (EE) embryos 7 DAP and late-stage (LE) embryos 25 DAP, representing the morphogenesis and maturation phases, respectively (Figure 4).

**Figure 4.**
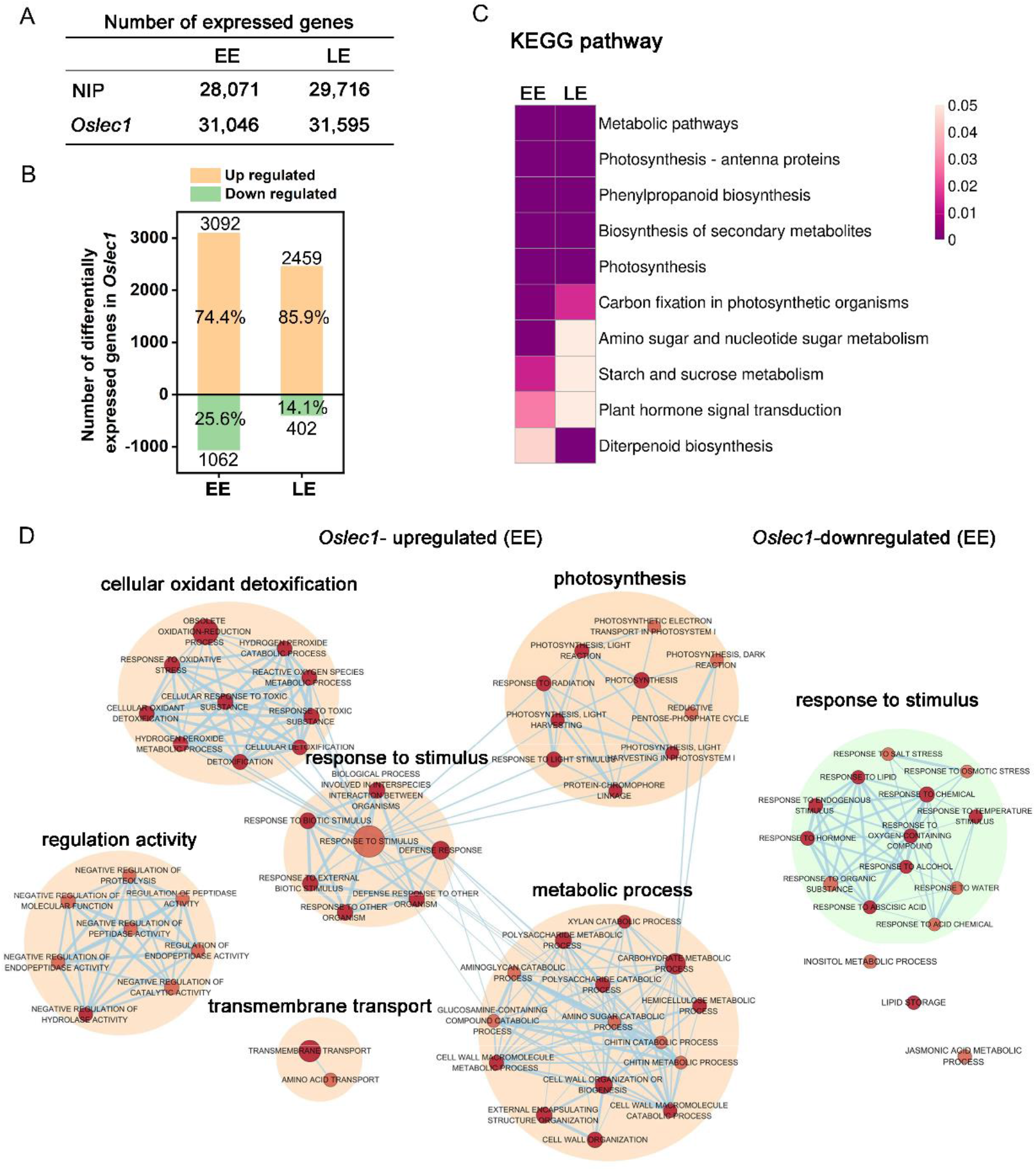
mRNA profiling of *Oslec1* mutant embryos at early (EE) and late (LE) development stages. **(A)** The number of differentially expressed genes detected in wild type and *Oslec1-1* embryos at EE (7 DAP) and LE (25 DAP) stages. **(B)** The number of upregulated and downregulated genes in *Oslec1-1* embryos compared with that in wild type plants. **(C)** KEGG pathway analysis of differentially expressed genes detected in wild type and *Oslec1-1* embryos at EE and LE stages. **(D)** GO term analysis of upregulated and downregulated genes in *Oslec1-1* embryos compared with those in the wild type ones.

Spearman correlation of 12 samples in the two stages showed good repeatability with a coefficient (R^2^) of above 0.99, and good heterogeneity with an R^2^ below 0.75 (Supplemental Figure 5). The number of all expressed genes detected in wild type and *lec1* mutant embryos at the two stages are shown in Fig. 3a. Compared with the wild type, there were 3092 and 2459 upregulated genes [p<0.05, log_2_ (Fold change)>=2] in *Oslec1-20* mutants at the two stages (EE and LE embryos), while there were only 1062 and 402 downregulated genes [p<0.05, log_2_ (Fold change) <= -2], respectively, (Figure 4B, Supplemental Table 1-4). The percentage of upregulated genes among the total number of differentially expressed genes (DEGs) (74.4 and 85.9%) was more than that of downregulated genes (25.6 and 14.1%), suggesting that OsLEC1 mutation activated more genes than it suppressed. Thus, OsLEC1 functioned mainly as a repressor in rice seed development.

Top Kyoto Encyclopaedia of Genes and Genomes (KEGG) pathways (p value<0.05) were similar in EE and LE stages (Figure 4C), including metabolic pathways, photosynthesis-antenna proteins, phenylpropanoid biosynthesis, biosynthesis of secondary metabolites, photosynthesis, and carbon fixation in photosynthetic organisms. KEGG terms related to amino sugar and nucleotide sugar metabolism, starch and sucrose metabolism, and plant hormone signal transduction were selectively enriched at the EE stage, and diterpenoid biosynthesis was preferentially enriched at the LE stage, as shown in Figure 4C and Supplemental Table 5. KEGG results suggested that OsLEC1 mutation affects many biological pathways involved in photosynthesis, biomacromolecule biosynthesis and metabolism, and hormone signal transduction in seed development.

To further explore functions of *OsLEC1* in the seed development, we identified all the GO terms significantly enriched (p value<0.01) in the biological progress of the two stages (Figure 4D, Supplemental Table 6-9). The network showed that upregulated genes were mainly enriched in GO terms involved in photosynthesis, cellular oxidant detoxification, response to stimulus, metabolic process and regulation activity, and downregulated genes were mainly enriched in GO terms involved in response to stimulus and biological process at EE and LE stages (Figure 4D, Supplemental Figure 6). The GO terms related to photosynthesis include photosynthesis; photosynthesis, light harvesting; photosynthesis, light reactions, consistent with the green-embryo phenotype of *Oslec1* mutants (Figure 2, 4D). GO terms involved in response to stimulus include response to water, ABA, hormones, and temperature stimulus, consistent with the drying intolerant and rapid germination phenotypes related to seed dormancy (Figure 1, 2, 4D). RNA-seq analysis demonstrated that *Oslec1* mutation played two roles in rice seed development: promoting photosynthesis-related pathways and inhibiting seed dormancy.

### *Oslec1* upregulates a series of genes involved in photosynthesis and photomorphogenesis

We designated genes regulated by OsLEC1 as those whose expression was at least four folds higher or lower [log2(Foldchange)>=2 or ≤ -2] in *Oslec1* mutants than that in wild type seeds at the same stage at a statistically significant level (p<0.05). Results showed that most genes involved in photosystem I (*OsPSA*s), photosystem II (*OsPSB*s), and light-harvesting complex (*OsLhcas* and *OsLhcbs*) were upregulated in the *Oslec1-1* mutant compared with the wild type (Figure. 5A). In addition, several DEGs involved in chloroplast development, as well as chlorophyll biosynthesis and degradation were also upregulated, such as *OsSGRL, OsPORA, OsPORB*, and *OsNYC4* (Figure 5A). The results suggested that OsLEC1 repressed many genes related to photosynthesis in rice seed development.

**Figure 5.**
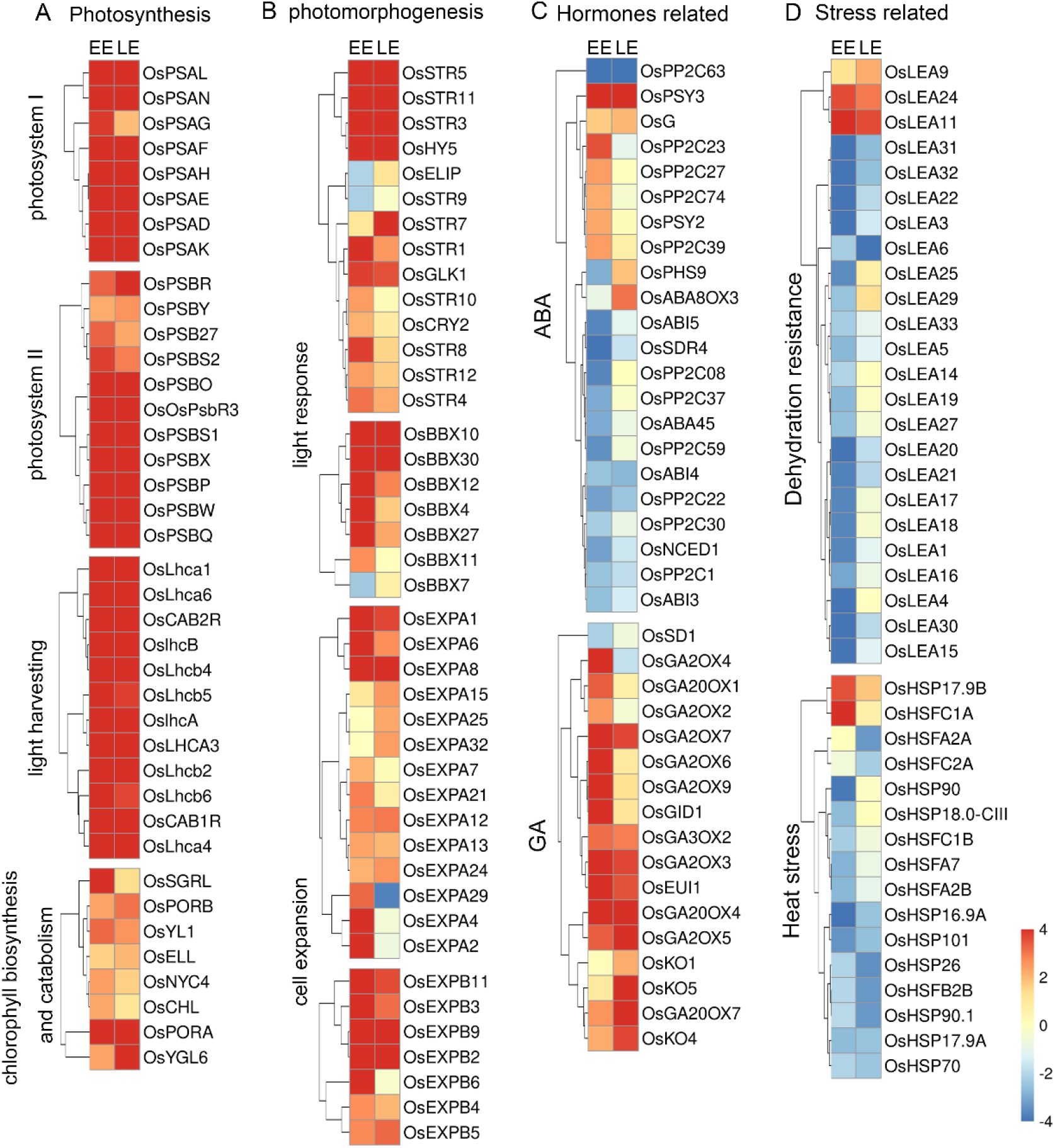
Heat maps of differentially expressed genes in *Oslec1-1* embryos. **(A)** Heat maps of genes involved in photosynthesis, **(B)** photomorphogenesis, **(C)** hormone pathways, **(D)** and stress responses, including dehydration resistance and heat stress response.

In addition, DEGs involved in photomorphogenesis, chloroplast development, and light response were analysed (Figure 5B). The results showed that *OsCRY2, OsHY5, OsBBX (4, 7, 10, 11, 12, 27, 30), OsEXPA (1, 2, 4, 6, 7, 8, 12, 13, 15, 21, 24, 25, 29, 32)*, and *OsEXPB (2, 3, 4, 5, 6, 9,11)* were significantly upregulated at least in one stage in the *Oslec1* mutant (Figure 5B, Supplemental Table 10). Ten members of the sulfurtransferase (Str) family (34), were upregulated in the *Oslec1* mutant at least in one stage, including *OsStr (1, 3*, 4, *5*, 7-*12)*. In addition, *OsGLK1* (35), was upregulated significantly at the EE and LE stages.

The above results demonstrated that *Oslec1* mutation upregulated a series of genes involved in photosynthesis and photomorphogenesis in the developing seeds.

### *Oslec1* regulates genes involved in diverse hormonal pathways and stress response contributing to seed dormancy

We identified the relative expression level of genes in the *Oslec1* mutant involved in hormone biosynthesis, signalling, and accumulation of six main hormones: ABA, GA, brassinosteroid (BR), zeatin, ethylene, and auxin (Figure 5C, Supplemental Figure 10). *OsPP2C (1, 08, 22, 30, 37, 59, 63), OsNCED1, OsSDR4*, and *OsABA45* involved in ABA biosynthesis, as well as *OsABI3, OsABI4*, and *OsABI5* involved in ABA signalling showed significant downregulation in the *Oslec1-1* mutant. Conversely, *OsKO (1, 4, 5), OsGA2ox (2, 3, 4, 5, 6, 7, 9), OsGA3ox2, OsGA20ox (1, 4, 7)*, and *OsGID1* involved in GA biosynthesis showed significant upregulation in the mutant (Figure 5C). In other hormonal pathways, several genes showed significant up-/downregulation. *OsCYP735A3* in zeatin biosynthesis and *OsHK2* in zeatin signalling, *OsBU1, OsSEPK1*, and *OsLAC15* in BR response, *OsACO (1, 2, 5)* in ethylene biosynthesis and *OsETR4* in ethylene signalling, *OsYUCCA (2, 6)* in auxin biosynthesis, *OsGH3-13* in auxin accumulation, *OsIAA (2, 12, 16)* and many *OsSAURs* in the auxin signalling pathway, were significantly upregulated (Supplemental Figure 7). Overall, the majority of the genes related to GA, zeatin, BR, ethylene, and auxin pathways were activated in the *Oslec1-1* mutant. In contrast, *Oslec1* mutation suppressed ABA biosynthesis and signalling in the developing seeds.

Stress response-related genes including *OsLEA*s, *OsHSP*s, and *OsHSF*s were also significantly changed in the *Oslec1-1* mutant (Figure 5D, Supplemental Table 11). Results showed that 21 *OsLEA*s were significantly downregulated, while only three were upregulated, such as *OsLEA (9, 11* and *24)*. Similarly, most *OsHSP* and *OsHSF* genes were downregulated. It is noteworthy that the expression levels of *OsLEA* genes showed increased downregulation at the EE stage than the LE stage, suggesting that *Oslec1* mutation in the regulation of *OsLEA*s mainly functioned in the early stage of seed development.

Overall, the transcriptome analysis of the *Oslec1* mutant suggested that OsLEC1 regulated genes involved in photosynthesis, photomorphogenesis, stress response, and diverse hormones pathways involved in seed dormancy.

### Genome-wide identification of OsLEC1 binding sites

To determine which processes are directly regulated by OsLEC1, we performed chromatin immunoprecipitation followed by high-throughput sequencing (ChIP-seq) in the callus of 35S:3xFLAG-OsLEC1 and the wild type. The peaks of 35S:3xFLAG-OsLEC1 were distributed mainly at the 5’-UTR site between the gene and 2 kb upstream, while the peaks of the wild type did not show a significantly enriched site and the normalised signal was significantly weaker than that of the 35S:3xFLAG-OsLEC1 (Supplemental Figure 8). In the 35S:3xFLAG-OsLEC1 group, 60.25% of the total 27452 peaks localised in the promoter region, including 51.96% within 1 kb of and 8.29% between 1-2 kb of the promoter (Figure 6A). The top enriched KEGG pathways (p<0.05) for these peak targets include amino sugar and nucleotide sugar metabolism (ko00520), butanoate metabolism (ko00650), N-glycan biosynthesis (ko00510), carotenoid biosynthesis (ko00906), and ribosome (ko03010) (Figure 6B, Supplemental Table 12).

**Figure 6.**
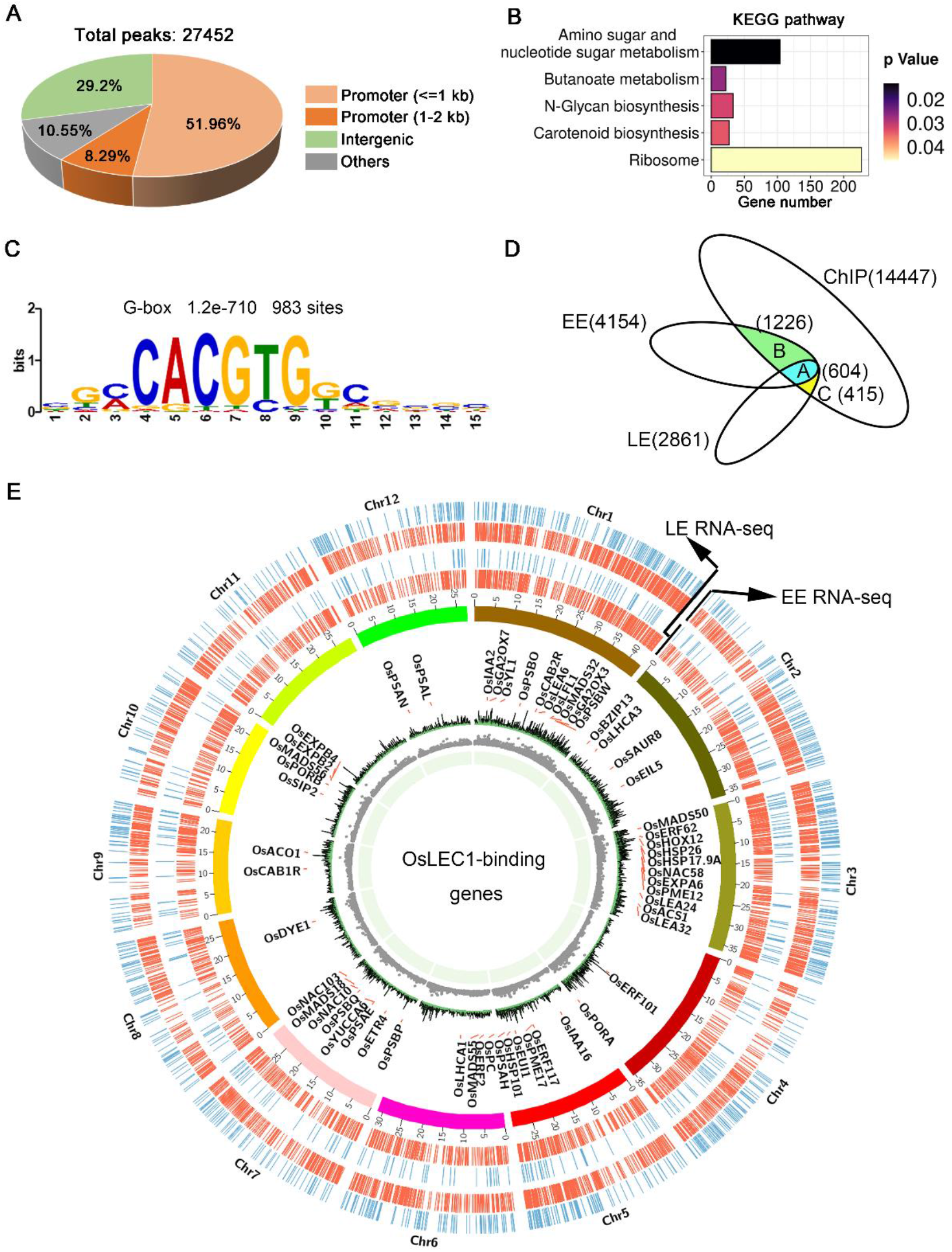
OsLEC1-binding genes detected using ChIP-seq. **(A)** The distribution of peaks on the rice genome according to ChIP-seq results. **(B)** KEGG pathway analysis of OsLEC1-binding genes. **(C)** DNA sequence motif G-box that enriched in LEC1-bound genomic regions 1 kb upstream of target genes, which were identified via de novo motif-discovery analyses. **(D)** Venn diagrams show the overlap between genes bound by OsLEC1 and differentially expressed genes in *Oslec1-1* embryos. Group A indicates 168 genes detected using ChIP-seq and RNA-seq at EE and LE stages; group B indicates 453 genes detected using ChIP-seq and RNA-seq at the EE stage; group C indicates 89 genes detected using ChIP-seq and RNA-seq at the LE stage. **(E)** Genome browser view of the chromosomal region showing enrichment of genomic regions bound by OsLEC1.

DNA sequence motifs that were enriched in LEC1-binding genomic regions 1 kb upstream of target genes were identified via de novo motif-discovery analyses (Figure 6C). These motifs most closely corresponded with several cis-regulatory elements including the G-box (CACGTG) or ABRE-like (C/G/T) ACGTG(G/T) (A/C), which were significantly overrepresented in OsLEC1 target genes with 983 binding sites. We did not detect a CCAAT DNA motif. A CIRCOS plot was constructed to identify and analyse similarities and differences between the OsLEC1 target genes according to ChIP-seq results with DEGs at EE and LE stages at the genome scale (Figure 6E). The gene density and number were larger at the EE stage compared with those at the LE stage. According to the annotation and the chromosomal distribution, OsLEC1 target genes were mapped for all 12 rice chromosomes with similar densities (Figure 6E). To determine the genes directly regulated by OsLEC1 in rice seed development, we analysed the overlapping OsLEC1-binding genes via ChIP-seq, using the DEGs detected in *Oslec1* embryos at EE and LE stages. There were 14447 unique target genes binding with OsLEC1, as well as 4,154 and 2,861 DEGs at EE and LE stage embryos of *Oslec1* mutants, respectively.

Venn diagram showed that OsLEC1 directly bound to 604 DEGs at EE and LE stages (Figure 6D, 6E), which were mainly involved in photosynthesis, chlorophyll biosynthesis, flowering, dehydration resistance, cell wall, heat stress, and hormonal pathways (GA, auxin, ethylene) (Supplemental Table 13-14). There were more target genes bound by OsLEC1 specifically at the EE stage (1226) than those at the LE stage (415) (Figure 6D), suggesting that OsLEC1 controlled several gene expression pathways in the early stage of seed development. At the EE stage, OsLEC1-binding genes were involved in the ABA pathway, dehydration resistance, lipid metabolic processes, heat stress, cell wall, and other hormonal pathways (auxin, cytokinin, ethylene). At the LE stage, OsLEC1-binding genes were involved in response to stress, seed development, JA, auxin-related and other transcriptional factors (Supplemental Table 13-14).

### OsLEC1 bound to DEGs in photosynthesis and seed maturation processes

According to the results above, among many DEGs involved in photosystem I (*OsPSA*s), photosystem II (*OsPSB*s), light-harvesting complex (*OsLhca*s), electron carrier (*OsPC*), and two genes encoding key enzymes in chlorophyll biosynthesis, *OsPORA* and *OsPORB*, were identified as OsLEC1-binding genes. These genes were upregulated (at least four folds compared with wild type) in *Oslec1* mutant embryos, contributing to photosynthesis (Figure 7A).

We identified *OsNCED1, OsSDR4, OsPP2C1*, and *OsABI3* as OsLEC1-binding genes, which were significantly downregulated in *Oslec1* embryos (Figure 7B). Downstream of *OsABI3*, 13 *OsLEA*s and 12 *OsHSF*s/*OsHSP90*.*1* were identified as OsLEC1-binding genes and were significantly downregulated in *Oslec1* embryos (Figure 7B).

**Figure 7.**
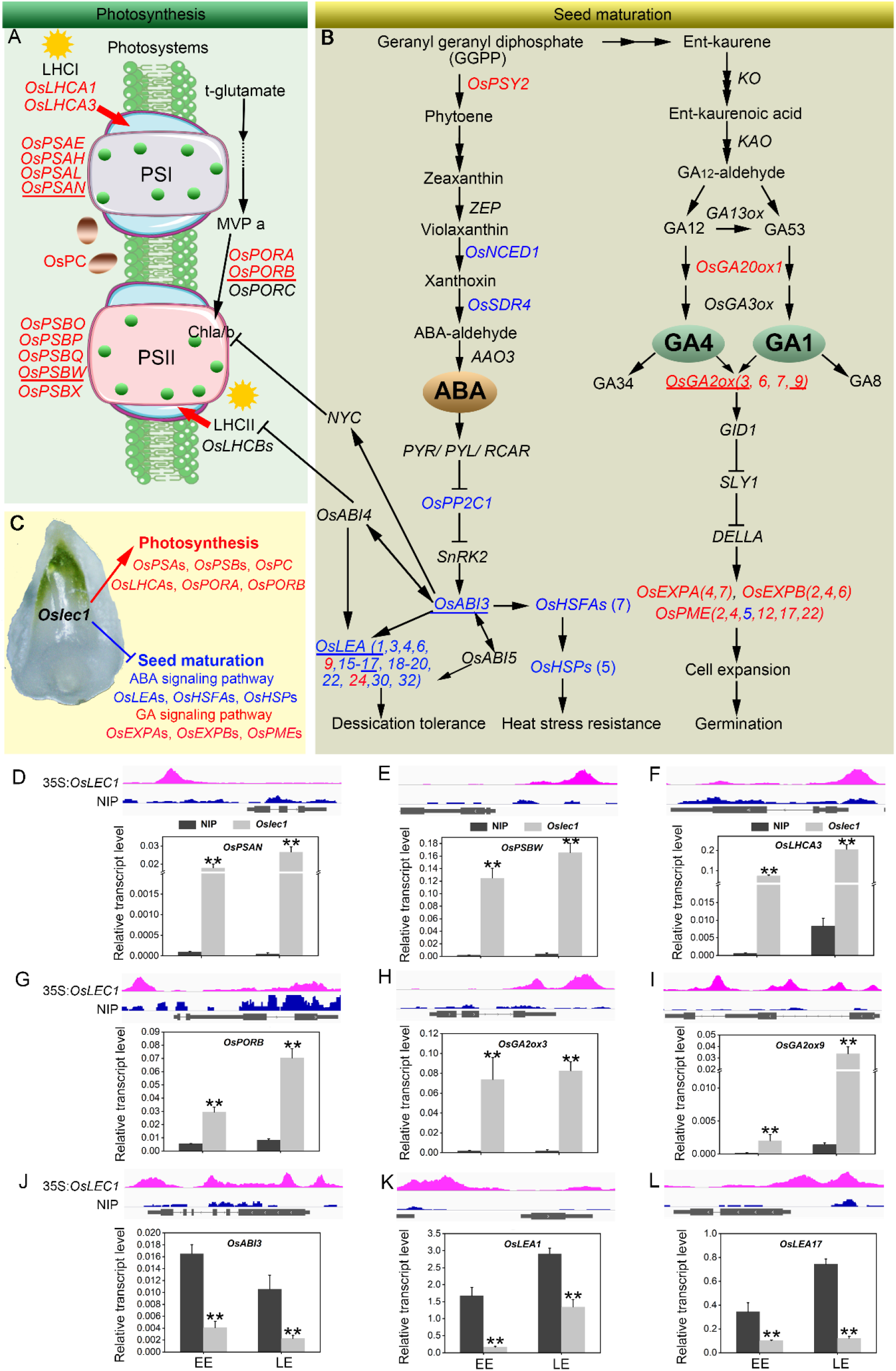
A proposed model of OsLEC1 action. **(A)** OsLEC1 directly binds and regulates genes involved in photosynthesis, including genes encoding proteins in PSI, PSII, LHCI, LHCII and OsPC, and key enzymes in chlorophyll a/b biosynthesis. **(B)** OsLEC1 directly binds and regulates genes involved in seed maturation, including genes in ABA/GA biosynthesis and signalling pathways. Genes in red indicate that they are upregulated in *Oslec1* embryos, and directly bound by OsLEC1; genes in blue indicate that they are downregulated in *Oslec1* embryos and directly bound by OsLEC1. **(C)** OsLEC1 induces desiccation tolerance, heat stress resistance, inhibition of early germination and seed dormancy, and inhibits photosynthesis in rice embryo development. OsLEC1 mutation results in the decrease of desiccation tolerance, heat stress resistance, inhibition of early germination and seed dormancy, as well as the activation of photosynthesis. **(D)** IGV screenshot of peak sites on genome sequences of OsLEC1-binding genes and qRT-PCR analysis of transcription levels of these genes in wild type and *Oslec1-1* embryos at EE and LE stages. These nine genes are all underlined in the above model.

We also found that *OsGA20ox1* and *OsGA2ox (3, 6, 7, 9)*, which are involved in GA biosynthesis and deactivation, were identified as OsLEC1-binding genes and were significantly upregulated in the EE- and LE-stage *Oslec1* embryos (Figure 7B). Cell expansion/cell wall-related genes *OsEXPAs, OsEXPBs*, and *OsPMEs*, downstream of the GA signalling pathway, were also identified as OsLEC1-binding and upregulated genes (Figure 7B).

Integrative genomics viewer (IGV) was used for the straightforward visualisation of OsLEC1-binding sites on nine target genes (Figure 7A, 7B). OsLEC1-binding sites of *OsPSAN, OsPSBW, OsLhca3, OsPORB, OsLEA1*, and *OsLEA17* were found on promoter regions, and those of *OsGA2ox3* and *OsABI3* were on the promoters and coding regions, while the binding site of *OsGA2ox9* was mainly on the coding region (Figure 7D-I).

Meanwhile, the expression levels of the nine genes in EE and LE stage embryos were detected using qRT-PCR. Results showed that *OsPSAN, OsPSAW, OsLhca3, OsPORB, OsGA2ox3*, and *OsGA2ox9* were upregulated in the *Oslec1* mutant, while *OsABI3, OsLEA1* and *OsLEA17* were significantly downregulated in early and late stages of seed development (Figure 7D-I). Overall, OsLEC1 mutation triggered photosynthesis but inhibited the seed maturation (Figure 7C).

## Discussion

LEC1 is an essential regulator of seed maturation (8, 13, 15, 18, 19, 36-38). The studies on LEC1 are summarised in Supplemental Figure 14. In the present study, we described the resulting phenotypes of the rice *Oslec1* mutant in seed development and illustrated an OsLEC1-binding/regulating gene network using a combination of RNA-seq and ChIP-seq data. We found that *Oslec1* mutation did not cause embryo lethality but showed an absence of dormancy, which caused death due to drying after seed harvesting (Figure 1). Chlorophyll accumulation in *Oslec1* embryos was triggered at a very early stage of seed development (Figure 2). OsLEC1 could directly bind and regulate many genes involved in photosynthesis, as well as ABA/GA biosynthesis and signalling pathway, contributing to dormancy (Figure 7). *Oslec1-1* and *Oslec1-2* seeds showed disrupted desiccation tolerance and failed to germinate after drying (Figure 1). Levels of major storage molecules were dramatically reduced in *Oslec1* mutant embryos (Figure 2). Notably, the shoot apices of *Oslec1* embryos were activated and possessed leaf primordia, whereas wild type embryonic shoot apices were inactive and did not initiate leaf development (Figure 2). Similar phenotypes were reported in Arabidopsis (12, 15, 19). A recent study showed that OsNF-YB7 (OsLEC1) complements the developmental defects of *lec1-1* in Arabidopsis when driven by the LEC1 native promoter (31). Therefore, OsLEC1 has a conservative function in seed maturation among dicots and monocots, which is required for the acquisition of desiccation tolerance and seed dormancy, storage compound accumulation, and inhibition of early germination (Figure 8Q).

**Figure 8.**
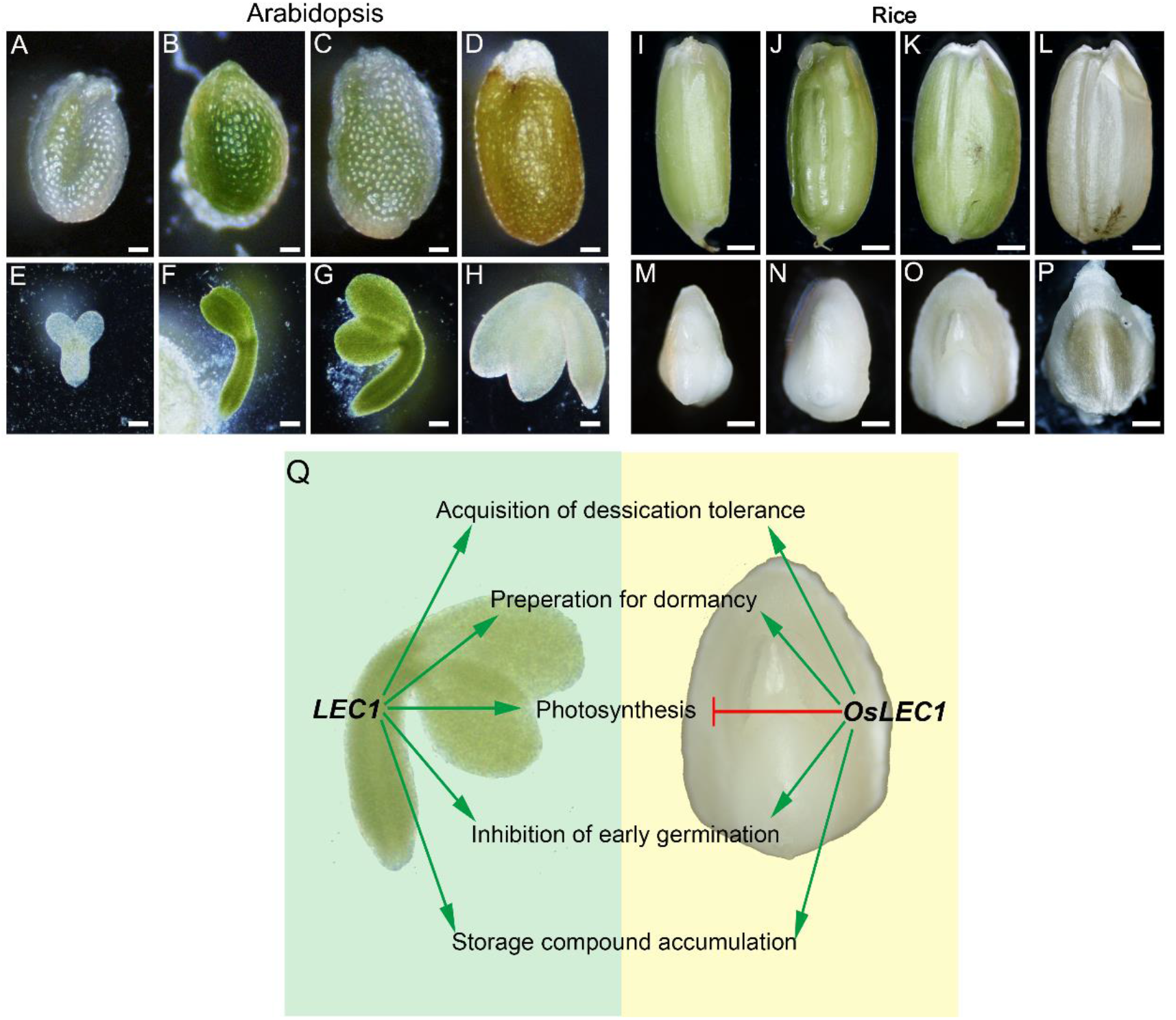
Developmental stages of Arabidopsis and rice seeds. **(A-D)** Complete Arabidopsis seeds in different developmental stages. Scale bars=200 μm. **(E-H)** Arabidopsis embryos in different developmental stages. Scale bars=50 μm. **(I-L)** Complete rice seeds in different developmental stages. Scale bars=500 μm. **(M-P)** Rice embryos in different developmental stages. Scale bars=50 μm. **(Q)** Comparison of LEC1 functions during seed development in rice and Arabidopsis. Arrowed green lines represent promotional effects, and a flat red line show inhibitory effects.

ABA and GA are major players in seed dormancy and germination regulation (39, 40). Some clues indicate the interaction of LEC1 and ABA/GA pathways. LEC1-induced embryonic differentiation in vegetative seedlings is strictly dependent on an elevated level of ABA (41). LEC1/L1L-[NF-YC2] activation depends on ABA-response elements (ABRE) present in the promoter of cruciferin C (CRC), which encodes a seed storage protein (42), and the ABRE-like motif (C/G/T)ACGTG(G/T)(A/C) was significantly overrepresented in LEC1 target genes (18). In addition, LEC1 interacts with the GA signalling repressor DELLA proteins. GA triggers the degradation of DELLAs to relieve their repression of LEC1, thus promoting auxin accumulation to facilitate embryonic development (43). Furthermore, LEC1 interacts with different combinations of AREB3, bZIP67, and ABI3 to control diverse biological pathways during soybean seed development. LUC assay results showed that *NCED1, AREB3, GA3ox1*, and *GA20ox2* are bound and directly regulated by LEC1 (32). We showed that OsLEC1 bound to and positively regulated *OsNCED1, OsSDR4, OsPP2C*s, and *OsABI3* in ABA biosynthesis and signalling but negatively regulated *GA2ox3* and *GA2ox9* in the GA pathway (Figure 7A, 7B), consistent with previous studies. Taken together, OsLEC1 plays a conservative role in controlling seed maturation through the regulation of ABA and GA pathways.

Owing to the non-photosynthetic nature of the rice seed, the rice embryo remains white during seed development and maturation (Figure 8A-H), different from the Arabidopsis seed which undergoes embryo greening (Figure 8I-P). Notably, *Oslec1* mutation could induce photosynthesis (Figure 7C). *Oslec1* embryos turned green 7 DAP (Figure 2), showing that photosynthesis had been activated in the very early stages of seed development. Compared with wild type plants, a series of photosynthesis-related genes were significantly upregulated in *Oslec1* embryos (Figure 5A, 5B), and many of them were also OsLEC1-binding targets (Figure 6E), showing a direct suppressive effect of OsLEC1 on photosynthesis (Figure 8Q). The results revealed that OsLEC1 is a key inhibitor of photosynthesis in rice embryo development.

In Arabidopsis, LEC1 interacts with PIF4 to coactivate genes involved in dark-induced hypocotyl elongation (44) and regulate LHCB genes through interaction with pirin, a protein that enhances TF binding in mammals (45). However, the role of LEC1 on photosynthesis was largely unknown. It was reported that maturing embryos of the *lec1* mutant display a paler green than wild type embryos, and most DEGs involved in photosynthesis and chloroplast biogenesis are downregulated in Arabidopsis *lec1* mutant (18). LUC assay showed that *PSBW* and PSBP-1 involved in photosynthesis system II is activated by LEC1 in soybean embryo cotyledon cells, and LEC1 may interact with different TFs to activate distinct gene sets (32). Thus, LEC1 could promote but is not completely required for photosynthesis and chloroplast biogenesis in Arabidopsis and soybean seed development (18, 32). There seems to be conflicting clues on our results about OsLEC1 blocking photosynthesis in rice seed maturation. In contrast, there were also a few photosynthesis-related genes upregulated in mature greening- and postmature greening-stage embryos of Arabidopsis *lec1* mutant according to previous RNA-seq results (18). Besides, cotyledons of the *lec1* mutant stay green in the late seed maturation, suggesting that the defect of *lec1* might act on chlorophyll degradation (31).

An explanation of the differences in the effect of LEC1 on photosynthesis in seed maturation between rice and Arabidopsis possibly lies in the different seed developing processes. Photosynthesis is activated sequentially during Arabidopsis embryo development (43, 46). The rice seed is non-photosynthetic, which has an endosperm for nutrient supply in the early stage of plant growth, and photosynthesis is completely absent in the embryo during the seed development. Thus, the activation of photosynthesis in *Oslec1* mutants was easily observed in rice seed maturation in the present study. *OsLEC1*, a monocot homologue of *LEC1*, blocks photosynthesis and chloroplast biogenesis in rice embryos. Evidence showed that *OsNF-YB7* (*OsLEC1*) expression could complement the developmental defects of Arabidopsis *lec1-1*, such as the morphology of cotyledons and the desiccation tolerance, but the cotyledons were green, suggesting that *OsLEC1* did not block photosynthesis in Arabidopsis embryo (31). This implies that other genes controlling photosynthesis together with *LEC1*, also control Arabidopsis embryo maturation and greening. In addition, differences in the conservative region of the LEC1 protein sequence also suggested a possible aspect of LEC1 functional differences between dicots and monocots (Supplemental Figure 1). Therefore, we speculate that the role of OsLEC1 in rice is not equal to that of LEC1 Arabidopsis in photosynthesis (Figure 8Q).

In conclusion, we demonstrated that OsLEC1 plays a crucial opposing effect on rice embryo photosynthesis to promote dormancy. Revealing the underlying mechanism of how OsLEC1 regulates seed dormancy would help to provide strategies for enhancing grain dormancy levels and prevent PHS of crop species. Breakthroughs regarding OsLEC1 regulatory mechanisms on photosynthesis would expand our understanding of the molecular network underlying photosynthesis, thus contributing to improving photosynthesis in agriculture.

## Methods

### Plant materials and growth conditions

Wild type ‘Nipponbare’ rice (*Oryza sativa* L. ssp. *japonica*) and Columbia-0 (Col-0), as the wild type Arabidopsis, were used. *Oslec1, pOsLEC1:GUS, 35S:OsLEC1* were all generated in the ‘Nipponbare’ background. Plants were grown in a greenhouse under a 16 h light, 30°C/8 h dark, 24°C cycle.

For callus induction, sterile seeds were placed in the callus induction medium (N6 basal medium supplemented with 10 μM 2,4-D, pH 5.8) and incubated in the growth chamber under a 16 h light, 28°C /8 h dark, 24°C cycle (47).

For the germination assay, rice seeds (dry or wet) were unshelled and soaked in distilled water at 30°C in the dark until they germinated. Protruded or radical seeds were considered germinated seeds. Then, uniformly germinated seeds were placed in Petri dishes (12 cm) covered by filter paper soaked with distilled water and grown at 28°C and 90% relative humidity under 16 h light/8 h dark conditions.

### Plasmid construction and rice transformation

*Oslec1* transgenic plants were generated using the CRISPR-Cas9 technology, according to a previously described method (48). We designed two single gRNAs that specifically targeted the protein-coding regions of *OsLEC1*. The two gRNAs were assembled into a single vector using the polycistronic-tRNA-gRNA (PTG) strategy. Two gRNAs ‘CTCTGGGCCATGAGCCGCCT’ and ‘CAGACCGTCAACTTCGAGCA’ were assembled into a single vector *pRGEB32* using the polycistronic-tRNA-gRNA (PTG) strategy to construct *OsLEC1-PRGEB32*.

To generate the *pOsLEC1:GUS*, a 2841 bp genomic sequence upstream the ATG start codon of *OsLEC1* (*LOC_Os02g49370*) was cloned from the rice genome, and inserted into the *pENTR/D-TOPO* vector (Invitrogen, Carlsbad, CA, USA), and subsequently into the destination vector *pHGWFS7*, which contained a β-glucuronidase (GUS) gene fusion created via LR Clonase (Thermo Fisher Scientific, Waltham, MA, USA) reactions.

For the ChIP assay, *35S:3*FLAG-OsLEC1* was constructed first by fusing a sequence encoding the 3*FLAG peptide sequence with the *OsLEC1* cDNA, and the fusion was then inserted into pCAMBIA 1300 using infusion cloning according to the manufacturer’s instructions. The vectors *OsLEC1-PRGEB32, pOsLEC1:GUS*, and *35S:3*FLAG-OsLEC1* were introduced into rice callus using the Agrobacterium tumefaciens-mediated co-cultivation approach through the EHA105 strain. Homozygous T3 seeds were used for further experiments.

For subcellular co-localisation of proteins, the coding sequence, not including the stop codon of OsLEC1, was cloned into the pUGW5 vector to generate the *35S:OsLEC1-GFP* via LR Clonase reactions according to the manufacturer’s instructions. The *35S:OsIAA1-mcherry* vector has been described previously (49). The primers used are listed in Supplementary Table 15.

### SEM, TEM and light microscopy

For SEM analysis, 5-DAP and 25-DAP embryos of NIP and *Oslec1* were fixed overnight at 4°C in 2.5% glutaraldehyde in phosphate buffer (0.1M, pH7.0), washed three times in the phosphate buffer (0.1M, pH 7.0) for 15 min at each step, then, postfixed with 1% OsO_4_ in phosphate buffer for 2 h and washed three times in phosphate buffer. The samples were dehydrated through an ethanol series (30, 50, 70, 80, 90, 95 and 100% ethanol, 15 min each), then transferred to absolute ethanol, and dehydrated in Hitachi Model HCP-2 critical point dryer (Hitachi, Tokyo, Japan). The dehydrated sample was coated with gold-palladium in Hitachi Model E-1010 ion sputter for 4-5 min and observed using the Hitachi Model SU-8010 SEM (Hitachi, Tokyo, Japan).

For TEM, the samples were fixed in 2.5% glutaraldehyde and postfixed with 1% OsO4, then, dehydrated through an ethanol series, similar to the pre-treatment of SEM samples. After dehydration, samples were placed into absolute acetone for 20 min, then, into 1:1 and 1:3 mixtures of absolute acetone and the final Spurr resin mixture for 1 and 3 h, respectively, then, to the final Spurr resin mixture overnight.

Samples were placed in an Eppendorf containing Spurr resin and heated to 70 °C for more than 9 h, then, sectioned in a LEICA EM UC7 ultratome. Sections were stained with uranyl acetate and alkaline lead citrate for 5 and 10 min, respectively, and observed using the Hitachi Model H-7650 TEM.

For light microscopy, the embedded samples were sectioned at 2–5 μm thickness using a rotary microtome (Leica, Wetzlar, Germany) and stained with basic fuchsin, then, observed using a Nikon C-C phase turret condenser (Nikon, Tokyo, Japan).

### Histochemical analysis of GUS activity and TTC staining

For GUS staining, *pOsLEC1:GUS* embryos were incubated in the X-Gluc solution overnight at 37°C, as described previously (50), and then, fixed in 2.5% glutaraldehyde overnight to prepare them for imaging or semi-thin sectioning.

For TTC staining, the seeds were split longitudinally and soaked in 1% (w/v) TTC solution at 37°C for 3 h as described previously (51).

### Subcellular localisation of OsLEC1

Rice protoplasts were prepared and transformed as previously described (52). Approximately 8 µg of the expression vector *35S:OsLEC1-GFP* was transferred into rice protoplasts, then, the protoplasts were cultured at room temperature in the dark for 16 h. Fluorescence signals were detected and photographed using an LSM710 NLO confocal laser scanning microscope (Zeiss, Mannheim, Germany). Excitation/emission wavelengths were 488 nm for GFP and 561/575– 630 nm for mCherry. Fluorescence signals were analysed using the Zen2009 (Carl Zeiss) software.

### RNA extraction and qRT-PCR analysis

Total RNA was extracted from *Oslec1* and wild type embryos using the TRIZOL reagent (Invitrogen, Carlsbad, CA, USA) and cDNA was synthesised from 1 µg of total RNA using ReverTra Ace™ qPCR RT Master Mix with gDNA Remover (TOYOBO, Osaka, Japan). qRT-PCR analysis was performed on a Mastercycler ep realplex system (Eppendorf, Hamburg, Germany) using LightCycler 480 SYBR Green Master (Roche, Indianapolis, USA). *OsUBQ5* was amplified and used as an internal standard to normalise the expression of tested genes. The primers of the examined genes are listed in Supplementary Table 15.

### RNA-seq analysis

We collected about 30 *Oslec1* and wild type embryos (7 and 25 DAP) in triple and extracted mRNAs to perform Illumina sequencing. Library construction and deep sequencing were carried out using the Illumina Hiseq 2500 (Biomarker Technologies, Beijing, China). The adapters and low-quality reads were removed using Trimmomatic (version 0.36) (53). Afterwards, the clean reads were mapped to the rice reference genome (Oryza_sativa. IRGSP-1.0.45) using Hisat2 (version 2.1.0) (54). Transcripts were assembled and merged using StringTie (version 1.3.4d) (55). The DEGs were identified using DESeq2 (version 1.26.0) (56), with a false discovery rate of < 0.05 and an absolute value of log2 (fold change) > 2.

The GO terms and KEGG enrichment analyses were performed using the online platform g:Profiler (https://biit.cs.ut.ee/gprofiler/gost) (57). The directed acyclic graphs of GO terms were constructed using the online platform AgriGO v2.0 (http://systemsbiology.cau.edu.cn/agriGOv2/) (58). The heat-map of the DEGs were drawn using the online platform Omicstudio (https://www.omicstudio.cn/login). Construction of the GO term network was performed using Cytoscape (59). The RNA-seq data have been deposited in the Gene Expression Omnibus (GEO) database, www.ncbi.nlm.nih.gov/geo (Accession No: GSE179838).

### ChIP and ChIP-seq analysis

ChIP was performed as previously described (60). About 3 g of the wild type and *35S:3*FLAG-OsLEC1* callus were collected and treated with 1% formaldehyde for protein-DNA cross-linking. After fixation, chromatin was sonicated with Diagenode Bioraptor to generate 200–1000 bp fragments. Chromatin was immunoprecipitated with anti-DDDDK monoclonal antibody (MBL, Beijing, China). Then, chromatin-antibody complexes were precipitated with anti-IgG paramagnetic beads (GE Healthcare, Uppsala, Sweden). After six washing steps, complexes were eluted and reverse-crosslinked. DNA fragments were sent to Genergy Biotechnology (Shanghai, China) for ChIP-seq.

ChIP-seq libraries were prepared using Ovation Ultralow Library Systems (Nugen) according to the manufacturer’s instructions. Sequencing reads of the ChIP-seq were aligned using Bowtie2 (61) against the Oryza sativa IRGSP-1.0 genome assembly, and only uniquely mapped sequencing reads were retained. MACS2 (62) was used to call peaks compared to the input using q_value_thresholds=0.01. The aligned reads with biological replicates were processed based on the irreproducibility discovery rate (IDR) (63). Peaks were then annotated using ChIPseeker (64). MEME-ChIP (65) suite was used to discover the DNA binding motif. Visualisation of peaks on genomic regions was achieved with IGV 2 (66). OsLEC1-binding peak positions according to the ChIP-seq results and the DEGs at EE and LE stages between the *Oslec1* mutant and the wild type plants were visualized using CIRCOS (67). The ChIP-seq data have been deposited in the Gene Expression Omnibus (GEO) database, www.ncbi.nlm.nih.gov/geo (Accession No: GSE179596).

### Protein sequences analysis

Protein sequences of the *LEC1* homologous genes in dicots and monocots were obtained from the National Centre for Biotechnology Information database (https://www.ncbi.nlm.nih.gov/). The phylogenetic tree of the protein sequences was constructed with MEGA (https://www.megasoftware.net/) using the NJ method with the following parameters: Poisson correction, complete deletion, and bootstrap (1000 replicates, random seed). Multiple sequence alignments of proteins were performed using the ALIGNMENT software (https://www.genome.jp/toolsbin/clustalw).

## ACKNOWLEDGMENTS

We thank Yunrong Wu, Zhejiang University, for the management of the greenhouse, and appreciate the kind help from Weilan Wang, Junying Li, Nianhang Rong, and Li Xie in the Bio-ultrastructure analysis Laboratory of the Analysis Centre of Agrobiology and Environmental Sciences, Zhejiang University. Moreover, we thank Hua Wang and Zijuan Li, CAS Centre for Excellence in Molecular Plant Sciences, for their help in the study. This research was supported by the grants from the National Natural Science Foundation of China (Grant no. 31971932, 31771776, 31771477), Science Foundation of Zhejiang Province (Grant no. LGN21C130006), China Agriculture Research System (CARS-05-05A), and China Postdoctoral Science Foundation (508000-X91806).

