## supplemental materials for "Rice LEAFY COTYLEDON1 hinders photosynthesis in the embryo development to promote seed dormancy"

**This PDF file includes:**

Supplementary text
Figures S1 to S9
Tables S1 to S15

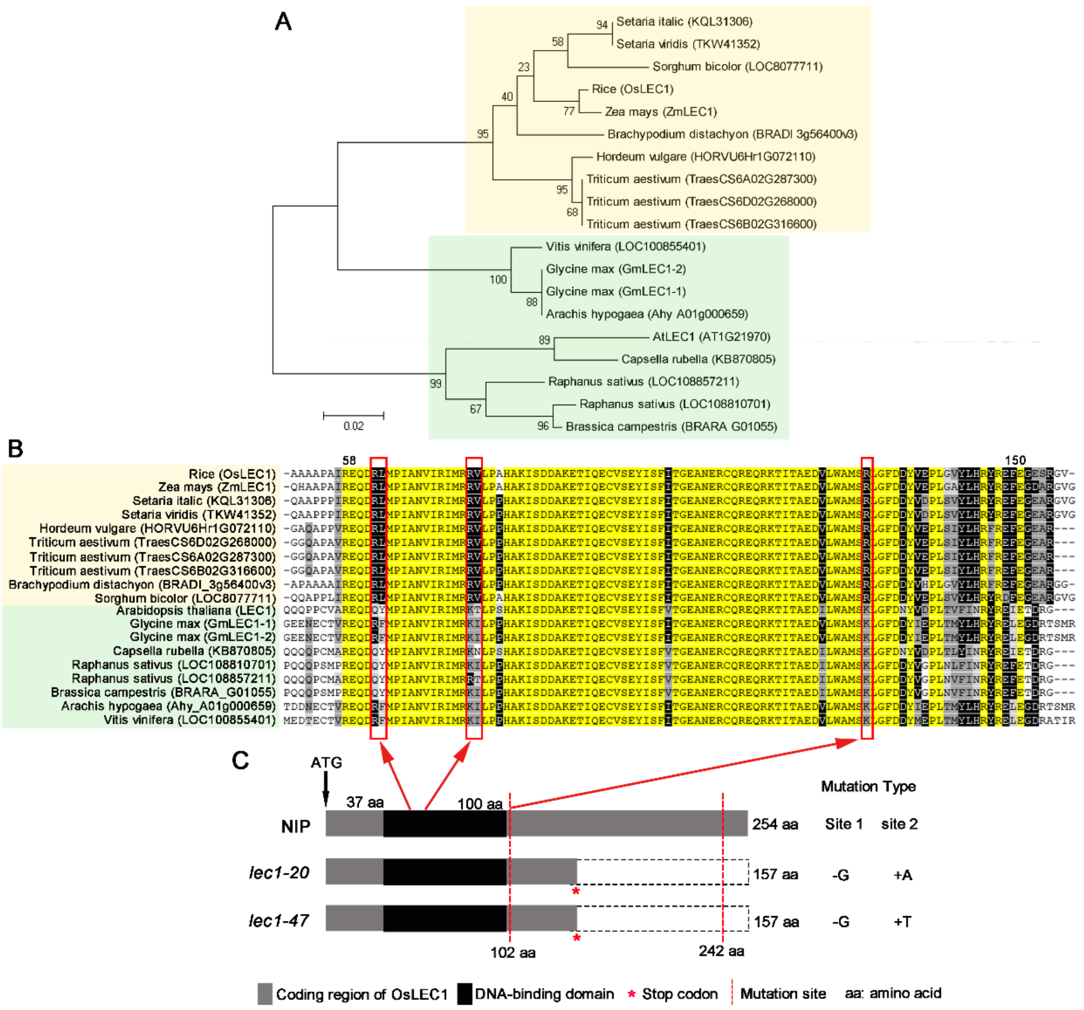


**Fig. S1.** Phylogenetic relationship and sequence alignment of LEC1 homologous proteins in dicots and monocots and the OsLEC1 protein sequence in the *Oslec1* mutant. **(A)** Phylogenetic analysis of LEC1 homologous proteins in dicots and monocots. **(B)** Sequence alignment of LEC1 homologous proteins. LEC1 naturally divided into three clades, indicating it diverged among the different clades. Multiple alignment results showed that LEC1 sequences harboured one conserved domain between amino acid (aa) 31–123 in rice, and the aa identity within the conserved domain reached up to 100%. While there were three sites in the conserved domain, it showed distinct differences between monocots and dicots: Arg 35-Leu 36 (RL); Arg 48-Val 49 (RV); Arg 102(R). The dicots and monocots are shown in light green and light yellow backgrounds, respectively. Identical residues are shown with white letters on a black background; conserved residues are shown with white letters on a grey background; a block of similar and weakly similar residues are shown with black letters on a light grey or white background. Red boxes indicate differential sequences in the conservative region between dicots and monocotyledons. **(C)** OsLEC1 protein sequence in the *Oslec1* mutant. The grey box with 254 aa represents the complete sequence of OsLEC1. The red asterisks indicate the premature translational termination site, the dotted blank boxes indicate the missing aa sequence and the red arrows indicate the differential sequences in the red boxes in C.

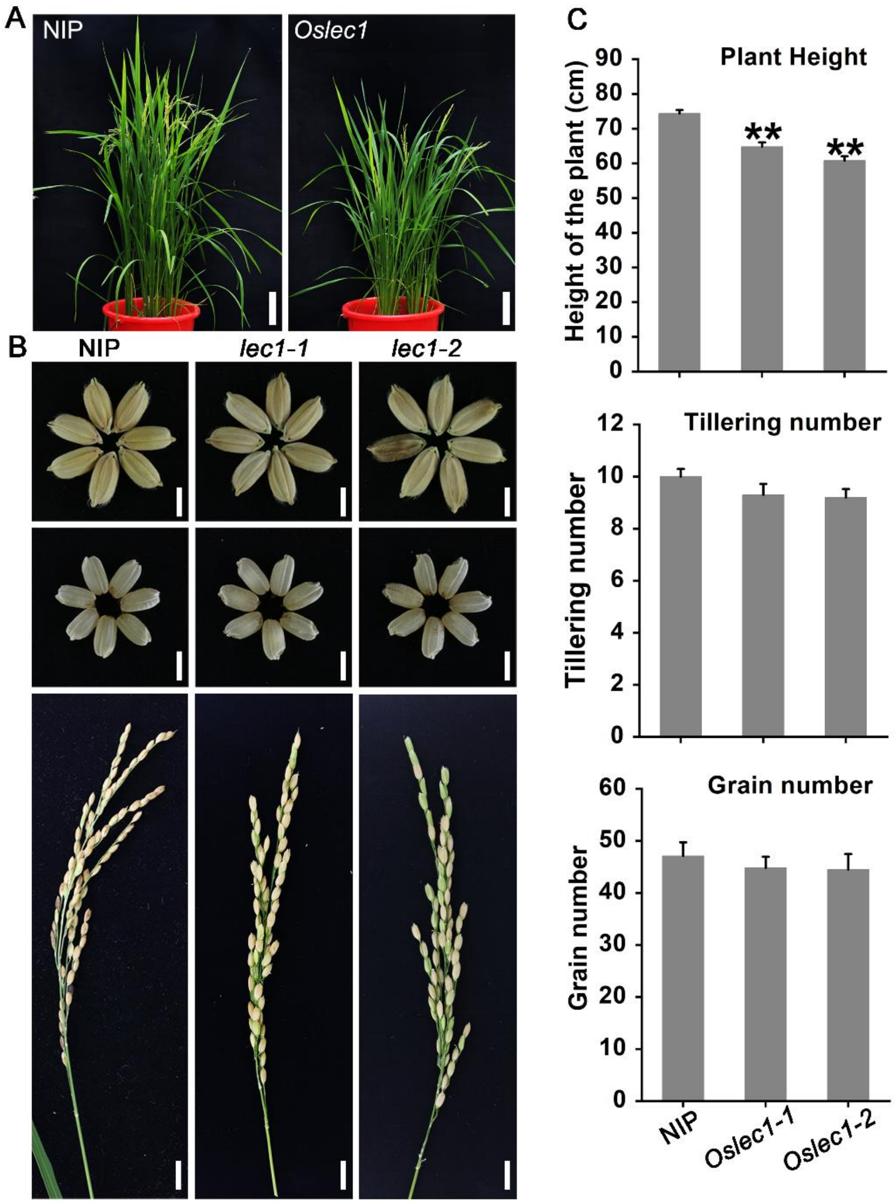


**Fig. S2.** Plant architecture and seed phenotype of the *Oslec1* mutant**. (A)** Architecture of the wild type plant and the *Oslec1* mutant about 3 months after germination. Scale bars=10 cm. **(B)** Mature seeds (Scale bars=5 mm) and panicles (Scale bars=1 cm) of the wild type plant and the *Oslec1* mutant. **(C)** Statistics of plant height, tillering number, and grain number/panicle of the wild type plant and the *Oslec1* mutant. Error bars indicate the SE of the mean; *P < 0.05; **P < 0.01 (Student's t-test), n>10.

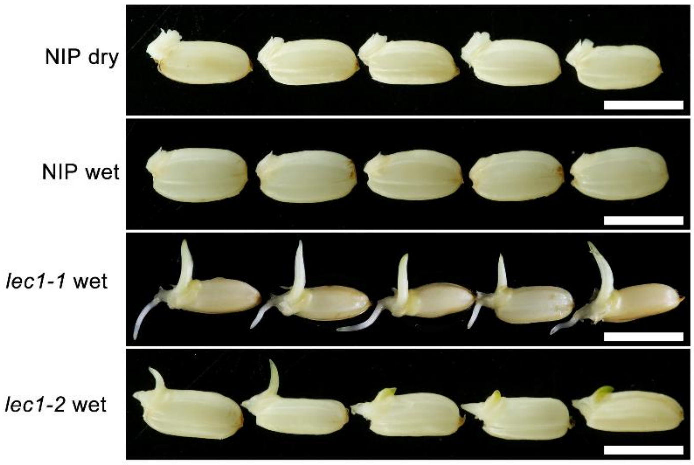


**Fig. S3.** Wild type and *Oslec1* mutant seeds 24 h after imbibition. Scale bars=5 mm.

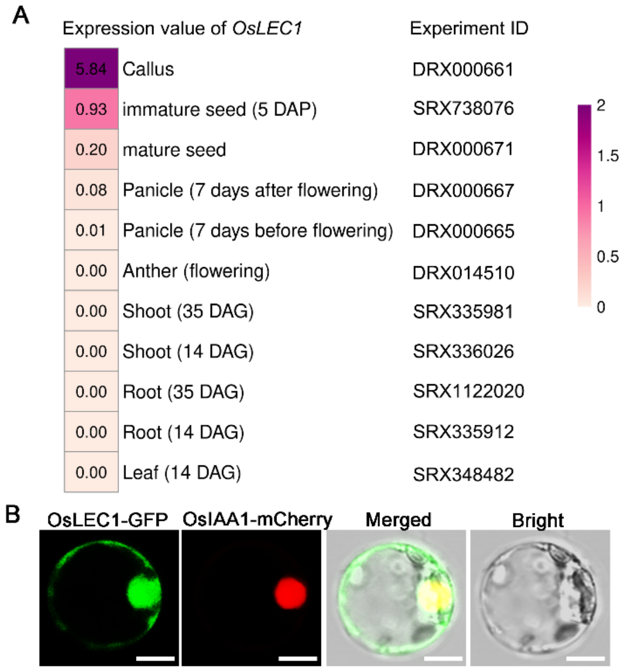


**Fig. S4.** Expression pattern of OsLEC1. **(A)** Expression of *OsLEC1* in different tissues and different stages. The data were obtained from http://http://expression.ic4r.org/. **(B)** Co-localization of OsLEC1-GFP with OsIAA1-mcherry. Both *35S:LEC1-GFP* and *35S: OsIAA1-mcherry* vectors were transformed into rice protoplasts. Fluorescence signals were observed under a confocal microscope. Scale bars=10 μm.

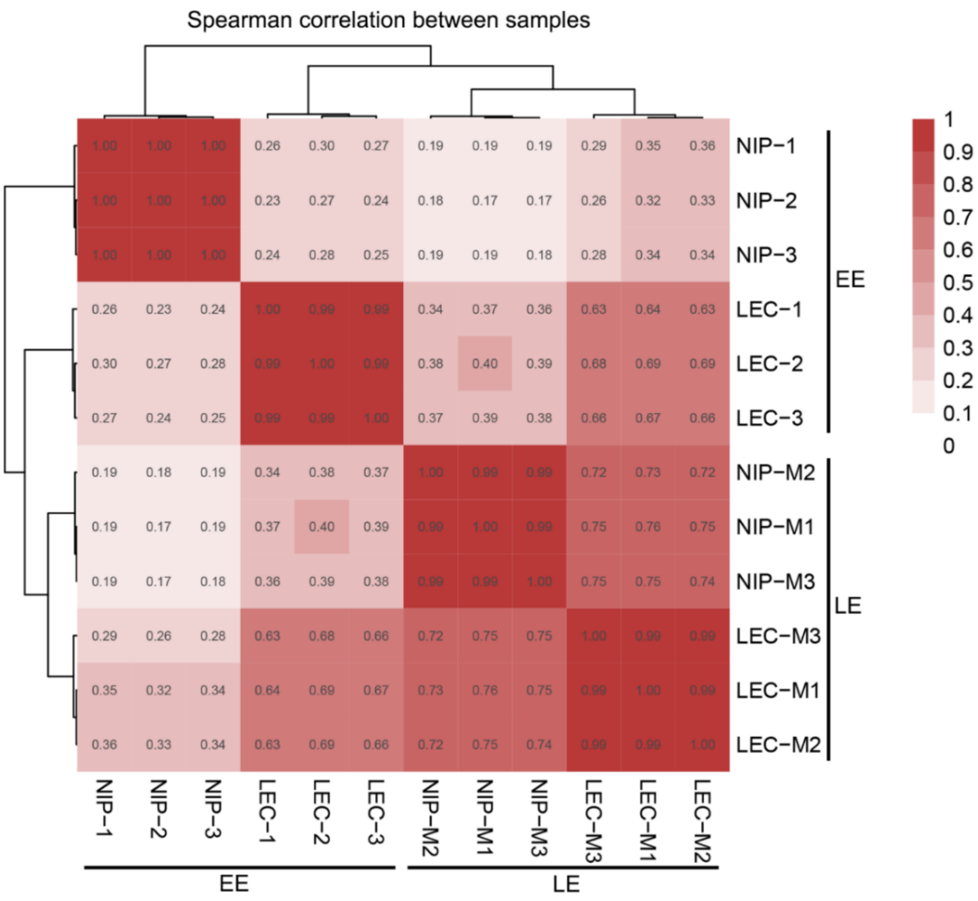


**Fig. S5.** Spearman correlation between 12 wild type and *Oslec1* mutant samples.

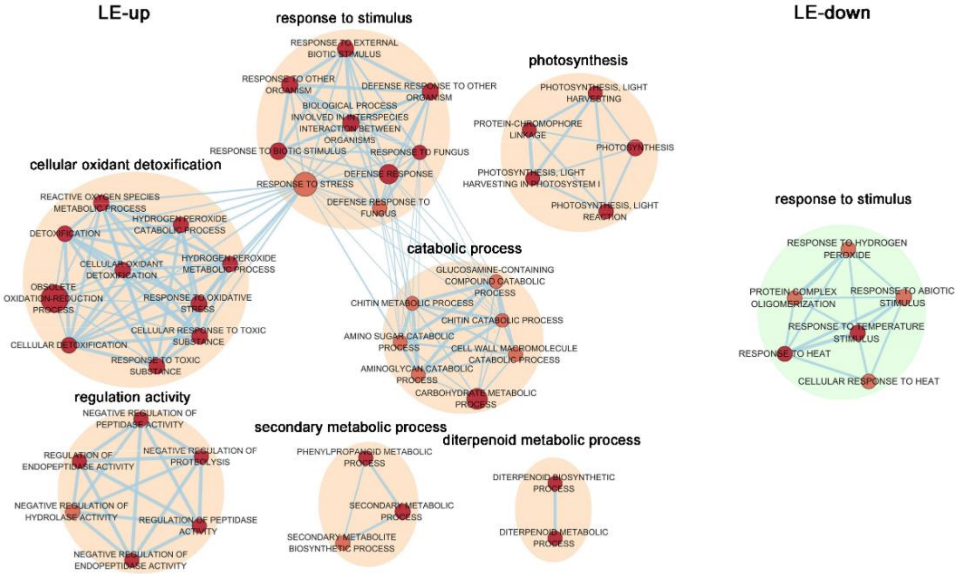


**Fig. S6.** GO term analysis of upregulated and downregulated genes in LE-stage *Oslec1* embryos.

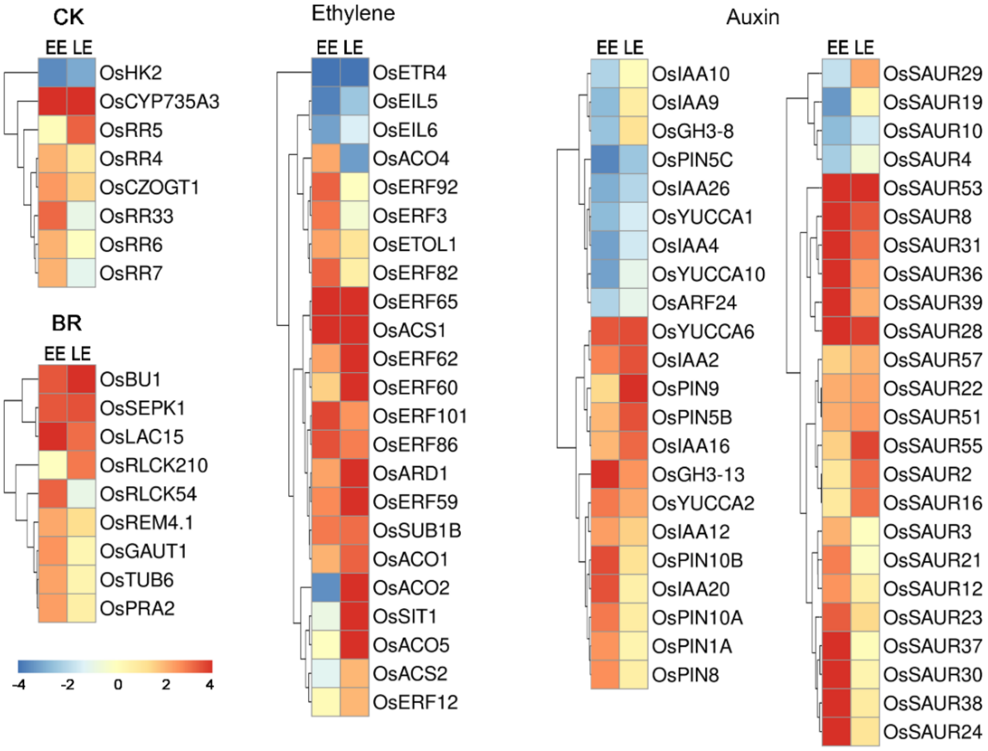


**Fig. S7.** Differentially-expressed genes in *Oslec1* mutant embryos in two stages involved in CK, BR, ethylene, and auxin pathways.

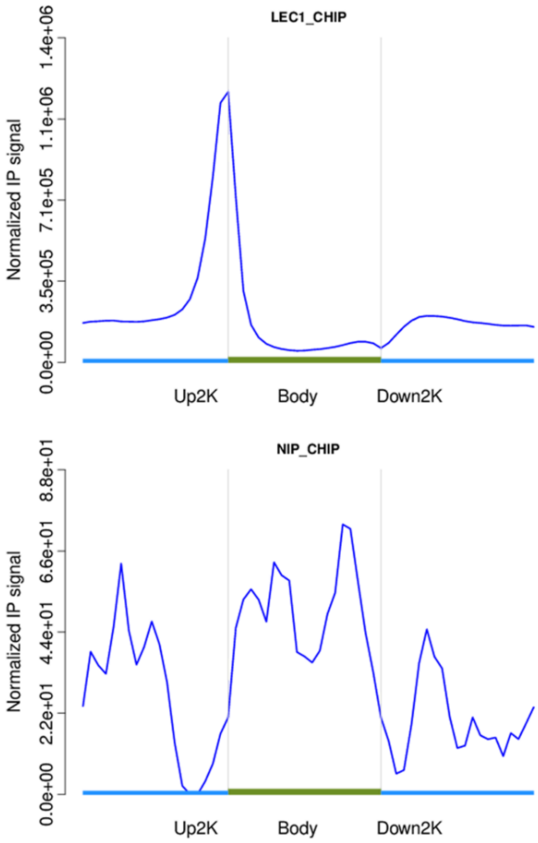


**Fig. S8.** Distribution of reads of ChIP-seq in gene bodies and upstream and downstream sequences.

| Species | Function | Genetic materials | Description in detail | References |
|---|---|---|---|---|
| Arabidopsis | Desiccation tolerance | *lec1* | Embryos of *lec1* mutants die of maturation drying at the end of seed development. | Meinke, 1992; Meinke *et al*., 1994; West *et al*., 1994 |
| Arabidopsis | Storage macromolecule accumulation | *lec1* | Storage protein and lipid accumulation are severely restricted in *lec1* mutants. | Meinke, 1992; Meinke *et al*., 1994; West *et al*., 1994 |
| Arabidopsis | Postgerminative seeding develepment | *lec1* | The shoot apices of *lec1* mutant embryos are activated and possess leaf primordia, whereas wild type embryonic shoot apices are inactive and do not initiate leaf development. | Meinke *et al*., 1994; West *et al*., 1994 |
| Arabidopsis | Embryo morphogenesis | *lec1*, *35S:LEC1* | *lec1* mutant embryo cotyledons, unlike wild type, undergo a heterochronic conversion in which they acquire leaf traits, such as trichomes on their adaxial surfaces and a cellular organization that is intermediate between cotyledons and leaves; trichome development is suppressed in plants overexpressing *LEC1*. | Meinke *et al*., 1994; West *et al*., 1994; Lotan *et al*., 1998; Huang *et al*., 2015 |
| Arabidopsis | Somatic embryogenesis | *lec1*, *35S:LEC1* | Ectopic expression of *LEC1* induces embryo development in vegetative organs. | Lotan *et al*., 1998; Gaj *et al*., 2005 |
| Kalanchoë daigremontiana | Asexual Reproduction | *pLEC1:C-KdLEC1* in K.daigremontiana backgroud | Expressing *KdLEC1* gene under the control of Arabidopsis *LEC1* promoter promotes viviparous leaf somatic embryos and thus enhances vegetative propagation in K. daigremontiana. | Garces *et al*., 2007; Garces *et al*., 2014 |
| carrot | Desiccation tolerance | *pLEC1:C-LEC1* in *lec1* background | Expressing carrot *C-LEC1* driven by the Arabidopsis *LEC1* promoter, could complement the viviparous and desiccation intolerant defects of Arabidopsis lec1-1 mutant. | Yazawa *et al*., 2004 |
| maize | Storage macromolecule accumulation, seed germination and leaf growth | EAP1:ZmLEC1, UBI:ZmLEC1 | Overexpression of maize *LEAFY COTYLEDON1* increases seed oil by as much as 48% but reduces seed germination and leaf growth in maize. | Shen *et al*., 2010 |
| rice | Development of leaves, panicles and spikelets | *pL::OsLEC1*, *pUN::OsLEC1* | Overexpression of *OsLEC1* resulted in abnormalities in the development of leaves, panicles and spikelets. | Zhang and Xue, 2013 |
| rice | Seed development | *Osnf-yb7* (*Oslec1*), *Osnf-yb9* | Defects of *OsNF-YB7* lead to lethality; Loss of *OsNF-YB9* function resulted in longer, narrower, and thinner seed. | Niu *et al*., 2021 |

**Fig. S9.** A summary of studies that report the functions of OsLEC1.
